# Activity of Tricyclic Pyrrolopyrimidine Gyrase B Inhibitor against *Mycobacterium abscessus*

**DOI:** 10.1101/2022.05.16.492225

**Authors:** Abdeldjalil Madani, Dereje A. Negatu, Abdellatif El Marrouni, Randy R. Miller, Christopher Boyce, Nicholas Murgolo, Christopher J. Bungard, Matthew D. Zimmerman, Véronique Dartois, Martin Gengenbacher, David B. Olsen, Thomas Dick

## Abstract

Tricyclic pyrrolopyrimidines (TPPs) are a new class of antibacterials inhibiting the ATPase of DNA gyrase. TPP8, a representative of this class, is active against *Mycobacterium abscessus in vitro*. Spontaneous TPP8 resistance mutations mapped to the ATPase domain of *M. abscessus* DNA gyrase and the compound inhibited DNA supercoiling activity of recombinant *M. abscessus* enzyme. Further profiling of TPP8 in macrophage and mouse infection studies demonstrated proof-of-concept activity against *M. abscessus ex vivo* and *in vivo*.

## MAIN TEXT

*Mycobacterium abscessus* causes difficult-to-cure lung disease (1). Multi-drug regimens are administered for months to years and typically contain an oral macrolide (clarithromycin, azithromycin), and intravenously administered amikacin, imipenem and / or cefoxitin or tigecycline. However, cure rates are unsatisfactory, and treatment refractory patients often undergo surgical lung resection. To further complicate treatment, the clinical utility of macrolides against *M. abscessus* is often limited by *erm41*-mediated inducible drug resistance (2). Given the poor performance of the current regimens, more efficacious drugs are needed. *M. abscessus* drug discovery efforts are hindered by extremely low hit rates in whole cell screens attempting to identify robust chemical matter starting points (3) (4).

*M. abscessus* is intrinsically resistant to many anti-tuberculosis (TB) antibiotics, including all first line drugs (5). Despite *M. abscessus* resistance to most approved anti-TB drugs, we found that compound collections of TB actives provide a good source for hit identification (6). Screening series of advanced TB actives against *M. abscessus* identified several compounds with *in vivo* activity, including inhibitors of RNA polymerase (7), ATP synthase (8), Leucyl-tRNA synthetase (9, 10), DNA gyrase (11) and DNA clamp DnaN (12). Expanding on this strategy, we asked whether the recently identified novel class of tricyclic pyrrolopyrimidines (TPPs (13)), targeting DNA gyrase in *Mycobacterium tuberculosis* and various other bacteria (14, 15) is active against *M. abscessus*.

DNA gyrase is a validated drug target in mycobacteria. This Type IIA DNA topoisomerase is an A_2_B_2_ heterotetrameric protein that regulates DNA topology (16). Unwinding of DNA during replication, transcription and recombination introduces positive supercoils into the DNA molecule that, left unaddressed impedes DNA function. This problem is resolved by DNA gyrase, which introduces negative supercoils into DNA. To do this, the enzyme generates a DNA double-strand break, passes a segment of DNA through the break, and subsequently reseals the DNA molecule (16). The fluoroquinolones target the cleavage-ligation active site of DNA gyrase formed by subunits A and B, creating stalled enzyme-DNA cleavage complexes (17).

Moxifloxacin is used effectively for the treatment of multi-drug resistant TB. However, the utility of this fluoroquinolone for treatment of *M. abscessus* infections is limited due to widespread intrinsic resistance (18). Recently, a novel benzimidazole (SPR719, Fig. 1A) entered early clinical development for mycobacterial lung diseases (19). Benzimidazoles target the ATPase domain of the DNA gyrase complex, located on its B subunits and required to drive the catalytic cycle (20), distinct from the fluoroquinolone binding site.

**Figure 1.**
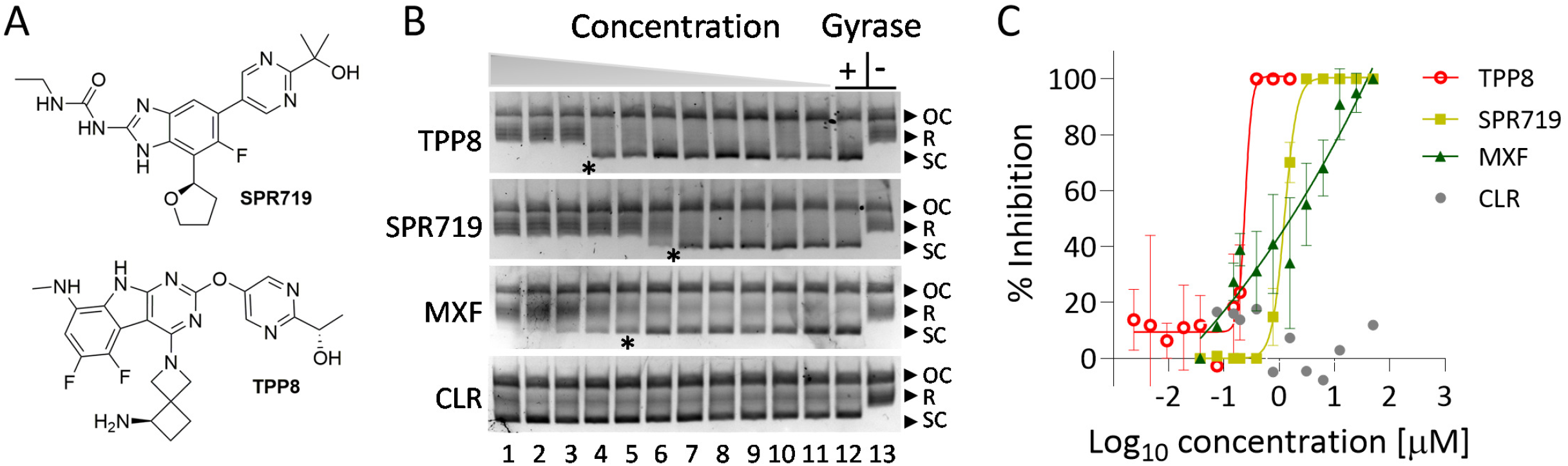
Structure and DNA gyrase inhibition activity of TPP8. **(A)** Structure of TPP8 and SPR719 (15, 20). **(B)** Effect of TPP8 and comparator compounds on the DNA supercoiling activity of recombinant *M. abscessus* ATCC19977 DNA gyrase. Relaxed pBR322 plasmid was used as substrate to measure the effect of compounds on the supercoiling activity of *M. abscessus* DNA gyrase as described (23). The conversion of relaxed (R) into supercoiled (SC) plasmid by DNA gyrase was visualized by agarose gel electrophoresis. OC: open circular plasmid. Lane 13, Gyrase -: reaction mix without added enzyme showing unaltered substrate. Lane 12, Gyrase +: reaction mix with added enzyme (without drug) showing conversion of relaxed plasmid into its supercoiled form. Lanes 1 to 11 show the effect of decreasing drug concentrations. The concentration ranges are as follows: TPP8: 1.5, 0.75, 0.37, 0.18, 0.09, 0.04, 0.02, 0.01, 0.005, 0.002, and 0.001 μM. SPR719, MXF and CLR: 50, 25,12.5, 6.25, 3.12, 1.56, 0.78, 0.39, 0.19, 0.09, and 0.04 μM. The experiments were repeated three times independently yielding similar results and a representative example is shown. **(C)** Quantitative inhibition of DNA gyrase supercoiling activity by TPP8 and comparator drugs. The bands obtained from the three experiments represented in **(B)** were quantified by the Invitrogen iBright™ FL1000 imaging system to determine half-maximal inhibitory concentrations (IC_50_) as described previously (23). Means and standard deviations are shown. TPP8 inhibited DNA gyrase with an IC_50_ of 0.3 μM. SPR719 and MXF inhibited the enzyme with an IC_50_ of 1 μM and 3 μM, respectively (23). IC_50_ derived from **(C)** are indicated by asterisks in **(B)**. CLR, included as negative control, did not affect the supercoiling activity of the enzyme.

Similar to SPR719, TPPs were shown to bind and inhibit the ATPase domain of the gyrase B subunit in *M. tuberculosis* (14). To determine whether this novel class of inhibitors is active against *M. abscessus,* the minimum inhibitory concentration (MIC) of a representative TPP compound (TPP8, compound #8 in (15) and Fig. 1A (21, 22), provided by Merck & Co., Inc., Kenilworth, NJ, USA was determined. Dose-response curves were established in Middlebrook 7H9 medium using the microbroth dilution method with OD_600_ as readout as described previously (23). TPP8 retained activity against reference strains from culture collections representing the subspecies of *M. abscessus*, including the type-strain *M. abscessus* subsp. *abcessus* ATCC19977, and a panel of clinical isolates, including *M. abscessus* subsp. *abscessus* K21, used in our mouse model of infection (Table 1). With growth inhibitory activity in the 0.02 to 0.2 μM range, TPP8 exhibited a markedly higher potency than SPR719 or moxifloxacin (Sigma-Aldrich), both showing MICs in the low micromolar range (Table 1). These results indicate that TPP8 is broadly active against the *M. abscessus* complex and displays potent antimycobacterial activity.

**Table 1.**
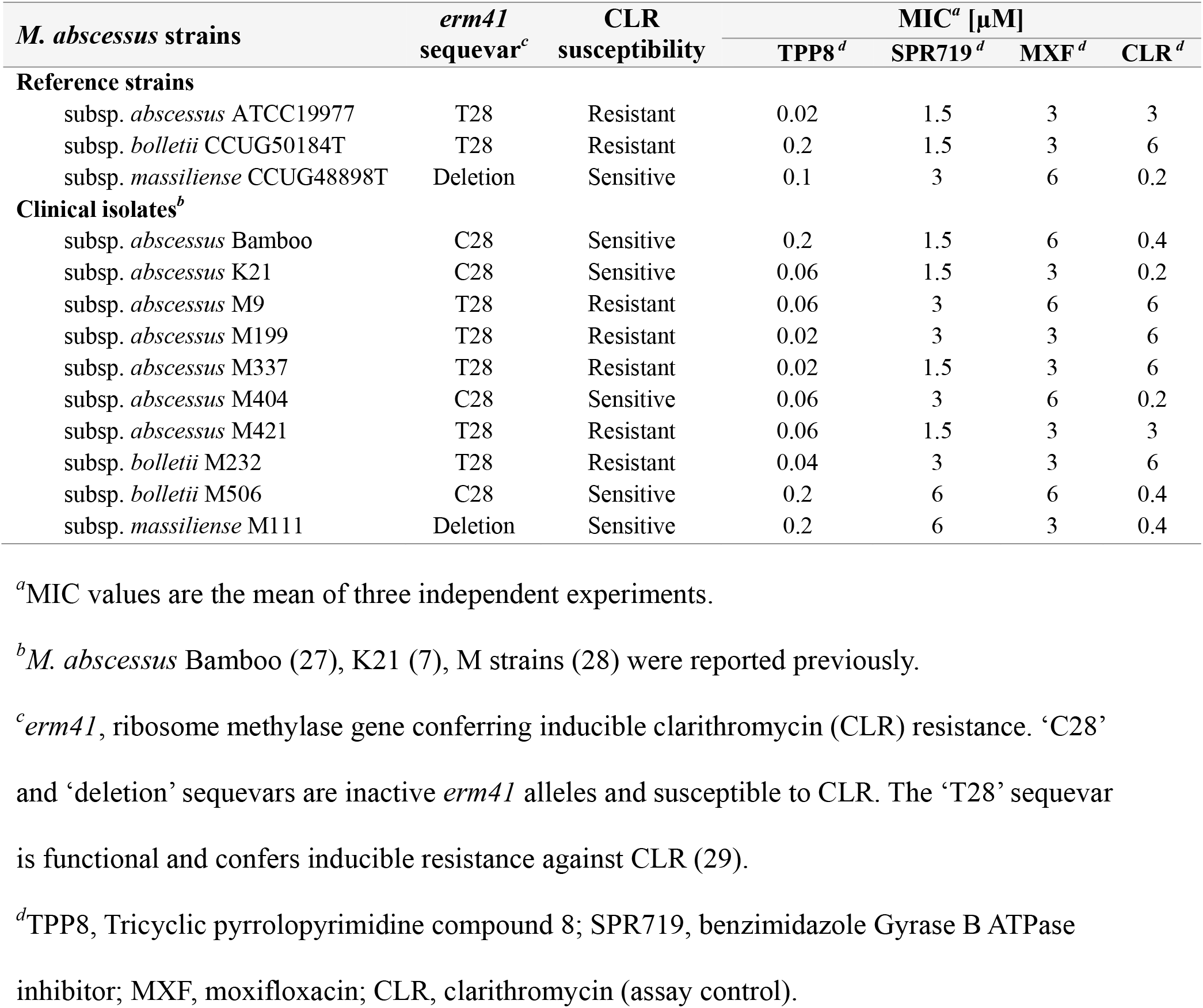
Activity of TPP8 against *M. abscessus* complex.

| *M. abscessus* strains | *erm41* sequevar<sup>c</sup> | CLR susceptibility | MIC<sup>a</sup> [μM] |  |  |  |
| --- | --- | --- | --- | --- | --- | --- |
|  |  |  | TPP8<sup>d</sup> | SPR719<sup>d</sup> | MXF<sup>d</sup> | CLR<sup>d</sup> |
| Reference strains |  |  |  |  |  |  |
| subsp. *abscessus* ATCC19977 | T28 | Resistant | 0.02 | 1.5 | 3 | 3 |
| subsp. *bolletii* CCUG50184T | T28 | Resistant | 0.2 | 1.5 | 3 | 6 |
| subsp. *massiliense* CCUG48898T | Deletion | Sensitive | 0.1 | 3 | 6 | 0.2 |
| Clinical isolates<sup>b</sup> |  |  |  |  |  |  |
| subsp. *abscessus* Bamboo | C28 | Sensitive | 0.2 | 1.5 | 6 | 0.4 |
| subsp. *abscessus* K21 | C28 | Sensitive | 0.06 | 1.5 | 3 | 0.2 |
| subsp. *abscessus* M9 | T28 | Resistant | 0.06 | 3 | 6 | 6 |
| subsp. *abscessus* M199 | T28 | Resistant | 0.02 | 3 | 3 | 6 |
| subsp. *abscessus* M337 | T28 | Resistant | 0.02 | 1.5 | 3 | 6 |
| subsp. *abscessus* M404 | C28 | Sensitive | 0.06 | 3 | 6 | 0.2 |
| subsp. *abscessus* M421 | T28 | Resistant | 0.06 | 1.5 | 3 | 3 |
| subsp. *bolletii* M232 | T28 | Resistant | 0.04 | 3 | 3 | 6 |
| subsp. *bolletii* M506 | C28 | Sensitive | 0.2 | 6 | 6 | 0.4 |
| subsp. *massiliense* M111 | Deletion | Sensitive | 0.2 | 6 | 3 | 0.4 |

<sup>a</sup>MIC values are the mean of three independent experiments.
<sup>b</sup>*M. abscessus* Bamboo (27), K21 (7), M strains (28) were reported previously.
<sup>c</sup>*erm41*, ribosome methylase gene conferring inducible clarithromycin (CLR) resistance. ‘C28’ and ‘deletion’ sequevars are inactive *erm41* alleles and susceptible to CLR. The ‘T28’ sequevar is functional and confers inducible resistance against CLR (29).
<sup>d</sup>TPP8, Tricyclic pyrrolopyrimidine compound 8; SPR719, benzimidazole Gyrase B ATPase inhibitor; MXF, moxifloxacin; CLR, clarithromycin (assay control).

To confirm that TPP8 exerts anti-*M. abscessus* whole cell activity via inhibition of gyrase B, spontaneous resistant mutants in *M. abscessus* ATCC19977 were selected on Middlebrook 7H10 agar as described previously (23). The agar MIC of TPP8 (lowest drug concentration that suppresses emergence of colonies when plating 10^4^ CFU on 7H10) was 0.64 μM as determined by the agar dilution method according to the CLSI protocol (24). To isolate spontaneous TPP8 resistant mutants, a total of 10^9^ CFU was plated on ten 30 mL agar plates containing 4x agar MIC, yielding one colony. TPP8 resistance was confirmed by re-streaking the colony on agar containing the same TTP8 concentration. The experiment was repeated once with an independently grown culture yielding a second TPP8 resistant *M. abscessus* strain. The broth MIC was similar for both mutants, 75-fold higher than the wild-type (Table 2). Susceptibility to moxifloxacin and clarithromycin (Sigma-Aldrich) was not affected, reducing the likelihood of a nonspecific mechanism of resistance (Table 2). Sanger sequencing of the gyrase B coding sequence, using primers GyrB-1 (GGCGTGGTGACGAGTTTAAAG), GyrB-2 (GAGATCTTCGAGACCACCACCTA), GyrB-3 (GCAAGAGTGCCACCGATATC) and GyrB-4 (GTAAGTACGACGGCACAACG) (Genewiz Inc.), showed that both resistant strains harbored a C506A (Thr169Asn) missense mutation, located in the ATPase domain (20) (Table 2). Interestingly, the same amino acid substitution in the *M. abscessus* gyrase B ATPase domain was previously shown to confer resistance to SPR719 (25). Indeed, cross resistance studies showed that the two TPP8 resistant *M. abscessus* ATCC19977 strains were resistant to SPR719 and that the previously isolated SPR719 resistant *M. abscessus* ATCC19977 strain harboring the C506A missense mutation (25) was resistant to TPP8 (Table 2). To confirm that the observed missense mutation in *gyrB* indeed causes resistance, the C506A allele of *gyrB* was overexperessed in wild-type *M. abscessus* ATTCC19977 using a custom synthesized (Genewiz Inc.) *pMN262-hsp60*-based expression system for *gyrBA* as described previously (23). The strain expressing the mutant enzyme showed resistance to both TPP8 and SPR719, confirming GyrB as the intracellular target (Table 2). To directly demonstrate that TPP8 inhibits *M. abscessus* DNA gyrase activity, *in vitro* DNA supercoiling inhibition studies were performed using recombinant *M. abscessus* enzyme and plasmid pBR322 (Inspiralis) as substrate, as described (23). The results demonstrate concentration-dependent enzyme inhibition by TPP8 (Fig. 1B,C). Consistent with the improved whole cell inhibitory potency of TPP8 compared to SPR719, the compound showed higher potency against the target with a half maximal inhibitory concentration (IC_50_) of 0.3 μM vs. 1 μM for SPR719. Together, these results provide genetic and biochemical evidence that TPP8 retained DNA gyrase B as its target in *M. abscessus*.

**Table 2.** Characterization of TPP8-resistant *M. abscessus* ATCC19977.

| *M. abscessus* ATCC19977 | MIC<sup>a</sup> (μM) |  |  |  | GyrB mutation |
| --- | --- | --- | --- | --- | --- |
|  | TPP8<sup>e</sup> | SPR719<sup>e</sup> | MXF<sup>e</sup> | CLR<sup>e</sup> |  |
| Wildtype | 0.02 | 1.5 | 3 | 3 | wt |
| TPP8<sup>R</sup>-1<sup>b</sup> | 1.5 | >25<sup>f</sup> | 3 | 1.5 | Thr169Asn |
| TPP8<sup>R</sup>-2<sup>b</sup> | 1.5 | >25<sup>f</sup> | 1.5 | 1.5 | Thr169Asn |
| SPR<sup>R</sup>-L1.2<sup>c</sup> | 3 | >25<sup>f</sup> | 3 | 2 | Thr169Asn |
| pMV262/ *hsp60 gyrB* *A<sup>d</sup> | 0.4 | 12.5 | 1.5 | 3 | wt |
| pMV262/*hsp60* empty<sup>d</sup> | 0.02 | 1.5 | 3 | 3 | wt |

<sup>a</sup> MIC values are the mean of three independent experiments.
<sup>b</sup> independently isolated TPP8 resistant mutant strains.
<sup>c</sup> SPR719 resistant mutant strain reported previously (25).
<sup>d</sup> wild-type strain expressing the DNA gyrase B\* and A subunits carried by pMV262 under control of *hsp60* (23) with Gyrase B\* harboring a Thr169Asn amino acid substitution.
‘pMV262/*hsp60* empty’, wild-type strain harboring the pMV262 expression system without gyrase genes inserted.
<sup>e</sup> TPP8, Tricyclic pyrrolopyrimidine compound 8; SPR719, benzimidazole Gyrase B ATPase inhibitor; MXF, moxifloxacin; CLR, clarithromycin (assay control).
<sup>f</sup> concentrations >25 μM could not be tested because of limited solubility of the compound (25).

To further characterize *in vitro* and *ex vivo* anti-*M. abscessus* activities of TPP8, kill experiments against *M. abscessus* ATCC19977 growing in Middlebrook 7H9 broth were performed and the inhibitory potency of TPP8 against bacteria growing intracellularly in infected THP-1 derived macrophages (ATCC TIB-202) was determined (26). TPP8 was largely bacteriostatic in broth culture (Fig. 2A) and inhibited growth of intracellular bacteria (Fig. 2B).

**Figure 2.**
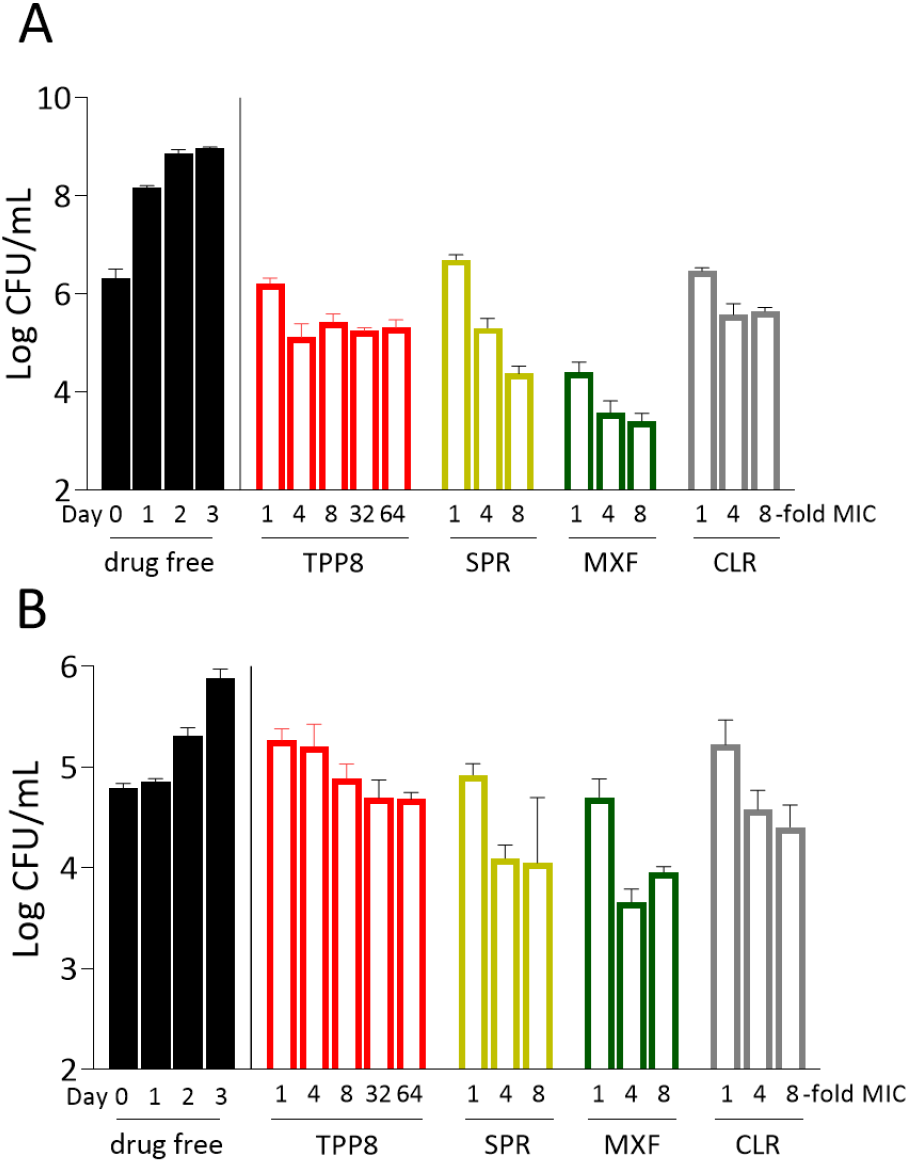
Activity of TPP8 against *M. abscessus* growing in broth and in THP-1 derived macrophages. **(A)** To determine whether TPP8 displays bactericidal activity *in vitro*, 1 mL cultures of *M. abscessus* ATCC19977 growing in Middlebrook 7H9 in tubes (11) were treated with MIC multiples of TPP8, SPR719, moxifloxacin (MXF), or clarithromycin (CLR). CFU were enumerated by plating samples on Middlebrook 7H10 agar. The growth kinetics of drug free controls are shown on the left, and the effects of TPP8 and comparators on CFU reduction are shown after 3 days of treatment. As MIC measured in tubes can be different from those measured in 96-well plates, tube MICs were measured and used as the baseline in these experiments (11). They were as follows (with MIC values shown in Table 1 and determined by the microbroth dilution method in parentheses): TPP8, 0.04 μM (0.02 μM); SPR719, 6 μM (1.5 μM); MXF, 6 μM (3 μM), CLR, 1.5 μM (3 μM). **(B)** To determine the activity against intracellular bacteria, THP-1 cells were prepared and differentiated into macrophages with phorbol-12-myristate-13-acetate for 24h, the resulting macrophages were infected with an MOI of 10 for 3h using *M. abscessus* ATCC19977 as described previously (26) and treated with the same concentration range of TPP8, SPR719, MXF, or CLR as in **(A)**. Intracellular CFU were enumerated by plating samples on agar Middlebrook 7H10 agar after 3 days of treatment. Experiments in **(A)** and **(B)** were carried out three times independently and the results are represented as mean values with standard deviations.

To determine whether the attractive *in vitro* and *ex vivo* activities of TPP8 translate into *in vivo* efficacy, an immunodeficient murine model developed by our group was utilized (7), in which mice are infected with the *M. abscessus* clinical isolate K21 (TPP8 MIC = 0.06 μM, Table 1) to generate a sustained infection resulting in a largely constant bacterial lung burden, thus allowing the effects of drugs to be evaluated (7). As TPP8 lacks robust oral bioavailability (15), the plasma concentration-time profile upon intraperitoneal administration in CD-1 mice (Charles River Laboratories) was determined. TPP8 plasma concentrations were measured by liquid chromatography-coupled tandem mass spectrometry. The *in vivo* pharmacokinetic analysis revealed that a dose of 25 mg/kg retains concentrations above the MIC of *M. abscessus* K21 for the 24h dosing interval (Fig. 3A). 8-week old female NOD.CB17-Prkdc^scid^/NCrCrl mice (NOD SCID; Charles River Laboratories) were infected by intranasal delivery of 10^6^ CFU as described previously (7). TPP8 was administered intraperitoneally once daily for 10 consecutive days at 25 and 12.5 mg/kg, starting one day post-infection. Two comparator agents were used in the efficacy study: the phosphate prodrug form of SPR719, SPR720 (20), administered orally at 100 mg/kg, and moxifloxacin administered orally at 200 mg/kg (11), the efficacious dose in TB mouse models (20). Clarithromycin as positive control was administered orally at 250mg/kg (11). All mice were euthanized 24h after the last dose, and bacterial load in the lungs and spleen was determined by plating serial dilutions of organ homogenates on Middlebrook 7H11 agar. All experiments involving live animals were approved by the Institutional Animal Care and Use Committee of the Center for Discovery and Innovation, Hackensack Meridian Health. As expected, treatment with vehicle alone did not affect the bacterial lung burden (‘D11 DF’, Fig.3B). Compared to the vehicle control, treatment with 25 mg/kg TPP8 reduced lung CFUs ~20-fold. The comparators SPR720 and moxifloxacin, and the positive control clarithromycin reduced the lung burden to a similar degree (Fig. 3B). CFU reduction in the spleen followed a similar pattern (Fig. 3B). Thus, TPP8 is efficacious in a mouse model of *M. abscessus* infection.

**Figure 3.**
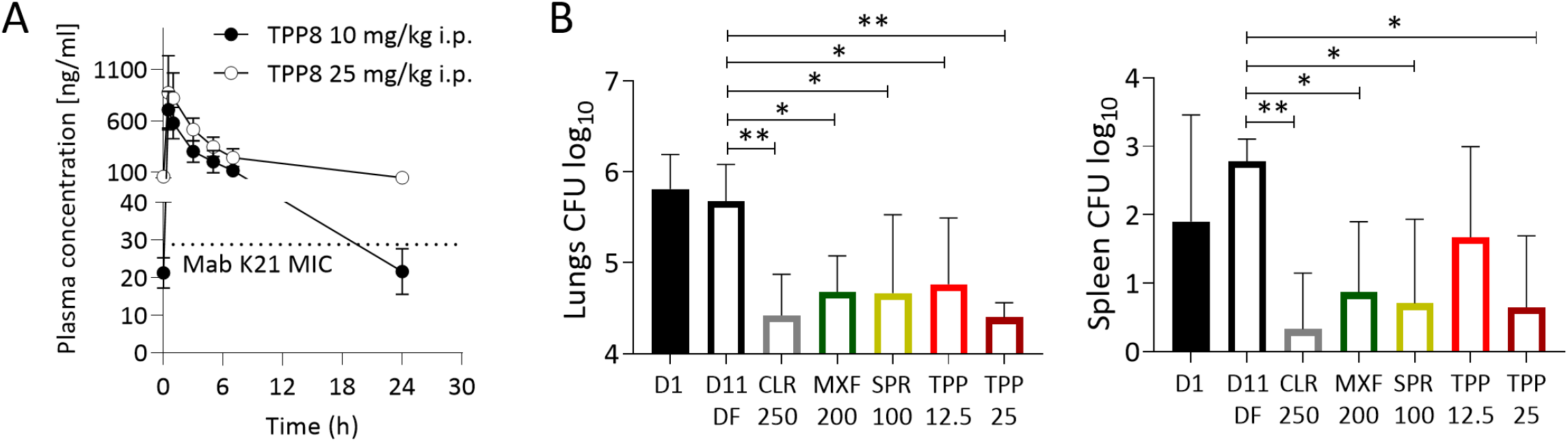
Pharmacokinetic profile and efficacy of TPP8 in mice. **(A)** Plasma concentration-time profile of TPP8 following a single intraperitoneal dose of 10 or 25 mg/kg in CD-1 mice. The MIC of TPP8 against *M. abscessus* K21 (Table 1), the strain used in our murine infection model, is indicated by a dotted line. **(B)** Efficacy of TPP8 and comparator compounds in a NOD SCID mouse model of *M. abscessus* K21 lung infection. Mouse lung and spleen CFU are shown one day after intranasal infection with *M. abscessus* K21 (D1), following daily intraperitoneal administration of 20% Solutol HS15 in PBS pH 7.4 (TPP8 vehicle) for 10 days (D11, DF: drug free), daily intraperitoneal administration of TPP8 (12.5 or 25 mg/kg), or daily oral administration of clarithromycin (CLR 250 mg/kg formulated in 0.5% carboxymethyl cellulose), moxifloxacin (MXF 200 mg/kg formulated in water) or SPR720 (SPR 100 mg/kg formulated in 0.5% methylcellulose) for 10 days. Mean and standard deviation are shown for each treatment group (n=6). Statistical significance of the results was analyzed by one-way analysis of variance (ANOVA) multi-comparison and Dunnett’s post-test: *, p<0.01; **, p<0.001. The experiment was carried out twice and one representative dataset is shown.

In conclusion, the tricyclic pyrrolopyrimidine TPP8 is active against *M. abscessus in vitro, ex vivo* and in a mouse model of infection and exerts its antimicrobial activity by inhibiting the B subunit of DNA gyrase. This work adds a new lead compound to the preclinical *M. abscessus* drug pipeline and provides an attractive chemical starting point for an optimization program aiming at improving oral bioavailability. The demonstration that yet another TB active displays anti-*M. abscessus* activity supports the strategy of exploiting chemical matter shown to be active against *M. tuberculosis* to accelerate *de novo* drug discovery for *M. abscessus*.

## ACKNOWLEDGMENTS

We are grateful to Wei Chang Huang (Taichung Veterans General Hospital, Taichung, Taiwan) for providing *M. abscessus* Bamboo, to Jeanette W.P. Teo (Department of Laboratory Medicine, National University Hospital, Singapore) for providing the collection of *M. abscessus* clinical M isolates, and to Sung Jae Shin (Department of Microbiology, Yonsei University College of Medicine, Seoul, South Korea) and Won-Jung Koh (Division of Pulmonary and Critical Care Medicine, Samsung Medical Center, Seoul, South Korea) for providing *M. abscessus* K21. We thank Wassihun Aragaw (Center for Discovery and Innovation, Hackensack Meridian Health, Nutley, New Jersey, USA) for providing the SPR719 resistant *M. abscessus* isolate. Research reported in this work was supported by the National Institute of Allergy and Infectious Diseases of the National Institutes of Health under Award Number R01AI132374. The content is solely the responsibility of the authors and does not necessarily represent the official views of the National Institutes of Health.

## AUTHOR CONTRIBUTIONS

Investigation: A.M., D.A.N., A.E.M., R.R.M., C.J.B., M.D.Z., M.G.; Materials: A.E.M.; Writing - Original Draft: A.M., D.A.N., T.D.; Writing - Review & Editing: all authors; Funding Acquisition: T.D., D.B.O.; Supervision: C.B., N.M., V.D., M.G., D.B.O., T.D.

